# RPL38 controls 60S ribosomal subunit homeostasis to regulate start-codon stringency and frame selection during *C9ORF72* RAN translation

**DOI:** 10.64898/2026.04.08.717149

**Authors:** Kohji Mori, Shiho Gotoh, Shizuko Kondo, Yuzo Fujino, Ryota Uozumi, Koujin Miura, Yuki Aoki, Shinji Tagami, Shoshin Akamine, Tesshin Miyamoto, Yoshitaka Nagai, Manabu Ikeda

**Author notes:** **Coressponding author:** Kohji Mori, M.D., Ph.D., Department of Psychiatry, Graduate School of Medicine, The University of Osaka, Yamadaoka 2-2, Suita, Osaka, 565-0871, Japan.

## Abstract

Translation initiation depends on stringent start-codon recognition, yet how ribosomal states modulate initiation stringency remains incompletely defined. Repeat-associated non-AUG (RAN) translation is a non-canonical initiation pathway that generates toxic dipeptide repeat proteins in *C9ORF72*-associated frontotemporal dementia and amyotrophic lateral sclerosis (FTD/ALS). Here, we performed a dual-luciferase reporter–based siRNA screen to identify factors that differentially regulate canonical AUG-initiated and *C9ORF72* RAN translation. We identified the 60S ribosomal protein RPL38 as a key determinant of translational output. RPL38 depletion selectively suppressed AUG-dependent translation while relatively preserving near-cognate–initiated RAN translation, resulting in a dose-dependent increase in the RAN-to-AUG translation ratio. Polysome profiling showed a selective reduction in large ribosomal subunit abundance without impaired subunit joining, consistent with defective 60S subunit homeostasis. Substitution of a near-cognate CUG start codon with AUG rendered translation sensitive to RPL38 depletion, supporting a role for ribosomal subunit availability in modulating initiation stringency. Graded RPL38 depletion altered frame usage within C9ORF72 repeat RNA. Consistent with these findings, RPL38 knockdown in the *Drosophila* eye enhanced GR-frame RAN translation while suppressing canonical AUG-dependent translation. Collectively, these findings identify 60S ribosomal subunit availability as a key determinant of start-codon stringency and frame selection in pathological *C9ORF72* RAN translation.

## Introduction

Translation initiation in eukaryotes is typically constrained by stringent recognition of AUG start codons during ribosomal scanning. However, initiation at near-cognate codons can occur under specific cellular conditions, including stress or altered translational states, yet the molecular determinants governing this shift in start-codon selection remain incompletely understood. In particular, how ribosome composition and subunit homeostasis influence initiation stringency and reading frame selection has not been fully elucidated.

Repeat-containing RNAs provide a unique context in which non-canonical initiation occurs with high efficiency and pathological consequences. Expanded nucleotide repeats, such as the GGGGCC (G4C2) repeat in *C9ORF72*, undergo repeat-associated non-AUG (RAN) translation to produce proteins from multiple reading frames in the absence of a canonical AUG start codon (1–4). In *C9ORF72*-associated frontotemporal dementia (FTD) and amyotrophic lateral sclerosis (ALS), this process generates toxic dipeptide repeat proteins (DPRs), including poly-GA, poly-GP, and poly-GR (2, 3), as well as antisense-derived DPRs (5–7), which accumulate in patient tissues and model systems. These DPRs exert toxicity through multiple mechanisms, including disruption of nucleocytoplasmic transport (8, 9), stress granule dynamics (10), DNA damage responses (11) , translation (12, 13), and proteostasis (14, 15). Notably, selective blockade of DPR synthesis without altering repeat RNA expression is sufficient to rescue multiple disease-associated phenotypes in vivo and in patient-derived neurons, underscoring the pathogenic importance of RAN translation (16).

Despite its importance, the molecular mechanisms that enable RAN translation remain incompletely understood (17). Mechanistically, C9ORF72 RAN translation largely exploits the canonical cap-dependent scanning pathway (18). Cryptic transcription and aberrant splicing generate m⁷G-capped repeat-containing transcripts that efficiently enter the cytoplasm and serve as templates for translation (19, 20). Cellular G4C2 repeat RNA abundance is tightly controlled by repeat RNA-binding proteins, such as hnRNPA3(11, 21–23), as well as by RNA decay pathways mediated by the RNA exosome (24, 25) . Ribosomes are recruited to the 5′ cap and initiate translation at near-cognate codons, including upstream CUG sites (18, 26–29), with occasional frameshifting enabling multi-frame translation (27, 30). Upstream open reading frames and local 5′ UTR context further modulate initiation efficiency (27, 31–33).

RAN translation is also tightly coupled to cellular stress responses. Activation of the integrated stress response (ISR), marked by phosphorylation of eIF2α, suppresses global cap-dependent translation while selectively enhancing RAN translation (18, 34). Structured repeat RNAs activate PKR, which functions as a major upstream regulator of ISR-dependent RAN translation (35). Canonical translation initiation factors further gate this process, as eIF5, eIF1, eIF1A, and eIF5B modulate near-cognate start-site selection during RAN initiation (29, 36, 37). In addition, repeat RNA secondary structures, including G-quadruplexes, influence ribosome engagement and elongation, and their remodeling by RNA helicases such as DHX36 or interactions with G-quadruplex–binding proteins dynamically modulate RAN translation efficiency (38–41). Small molecules that stabilize G-quadruplexes can stall ribosomes on repeat RNA and suppress elongation (42–44).

In parallel, accumulating evidence supports the concept of ribosomal heterogeneity, in which variations in ribosomal protein composition or subunit availability can bias translation toward specific subsets of mRNAs or alternative initiation mechanisms (45, 46). Ribosomal proteins are increasingly recognized as active regulators of translational specificity, influencing IRES-dependent initiation, uORF translation, and stress-responsive protein synthesis (47, 48). Consistent with this view, the small ribosomal subunit protein RPS25 has been shown to be selectively required for efficient RAN translation while being largely dispensable for canonical AUG-initiated translation (49).

Despite these advances, whether and how ribosomal subunit homeostasis directly influences start-codon selection and frame usage on C9ORF72 repeat RNA remains unclear.

Here, we systematically investigated the role of ribosomal and translation-associated factors in regulating canonical and non-canonical translation using a dual-reporter system that enables simultaneous monitoring of AUG-initiated and near-cognate–initiated translation. Through an siRNA-based screen, we identify the 60S ribosomal protein RPL38 as a key determinant of translational output. We show that perturbation of 60S ribosomal subunit homeostasis selectively impairs AUG-dependent translation while preserving RAN translation, resulting in a shift in translational balance and frame selection on G4C2 repeat RNA. These findings establish ribosomal subunit availability as a critical regulator of start-codon stringency and provide mechanistic insight into how ribosome composition shapes pathological non-canonical translation in *C9ORF72*-associated FTD/ALS.

## Results

### Dual-luciferase reporter screen to identify factors differentially affecting RAN and AUG translation

To monitor *C9ORF72* GGGGCC repeat RAN translation in three reading frames together with conventional AUG dependent translation, we developed dual luciferase reporter expression system (Figure 1A). Among the three RAN translation reporters, only poly-GA was detected by western blot, while poly-GP and poly-GR were undetectable (Figure 1B). To evaluate the relative luciferase activities of the three RAN translation reporters, an AUG-dependent Firefly luciferase (Fluc) construct was co-transfected with each reporter, and luciferase activities were subsequently quantified. In agreement with the western blot analysis and previous reports (2, 22) , poly-GA exhibited the highest activity, followed by poly-GR and poly-GP (Figure 1C, D). Next, we performed a siRNA-mediated knockdown screen of 24 genes involved in translation, including initiation factors, ribosomal components, and ribosome quality control, using the dual luciferase reporter system (Figure 1E). Most of the 24 genes targeted in this study were previously identified in CRISPR-based screens as candidate factors associated with altered RAN translation (38) . Using the poly-GA RAN translation reporter and AUG Fluc reporter in the primary screen, seven candidate genes that significantly alter RAN/AUG translation ratio were identified. Of these, knockdown of six genes (*RPS4X*, *RPL38*, *EIF3B*, *EIF3K*, *RACK1*, *GSPT1*) preferentially promoted poly-GA RAN translation relative to AUG translation (Figure 1F). Additional screening using the poly-GR RAN translation reporter, knockdown of *RPL38* or *RPS4X* again preferentially promoted poly-GR RAN translation relative to AUG translation (Figure 1G). Together, these results suggest loss of these candidate proteins may selectively alter the ratio of RAN translation/AUG translation.

**Figure 1.**
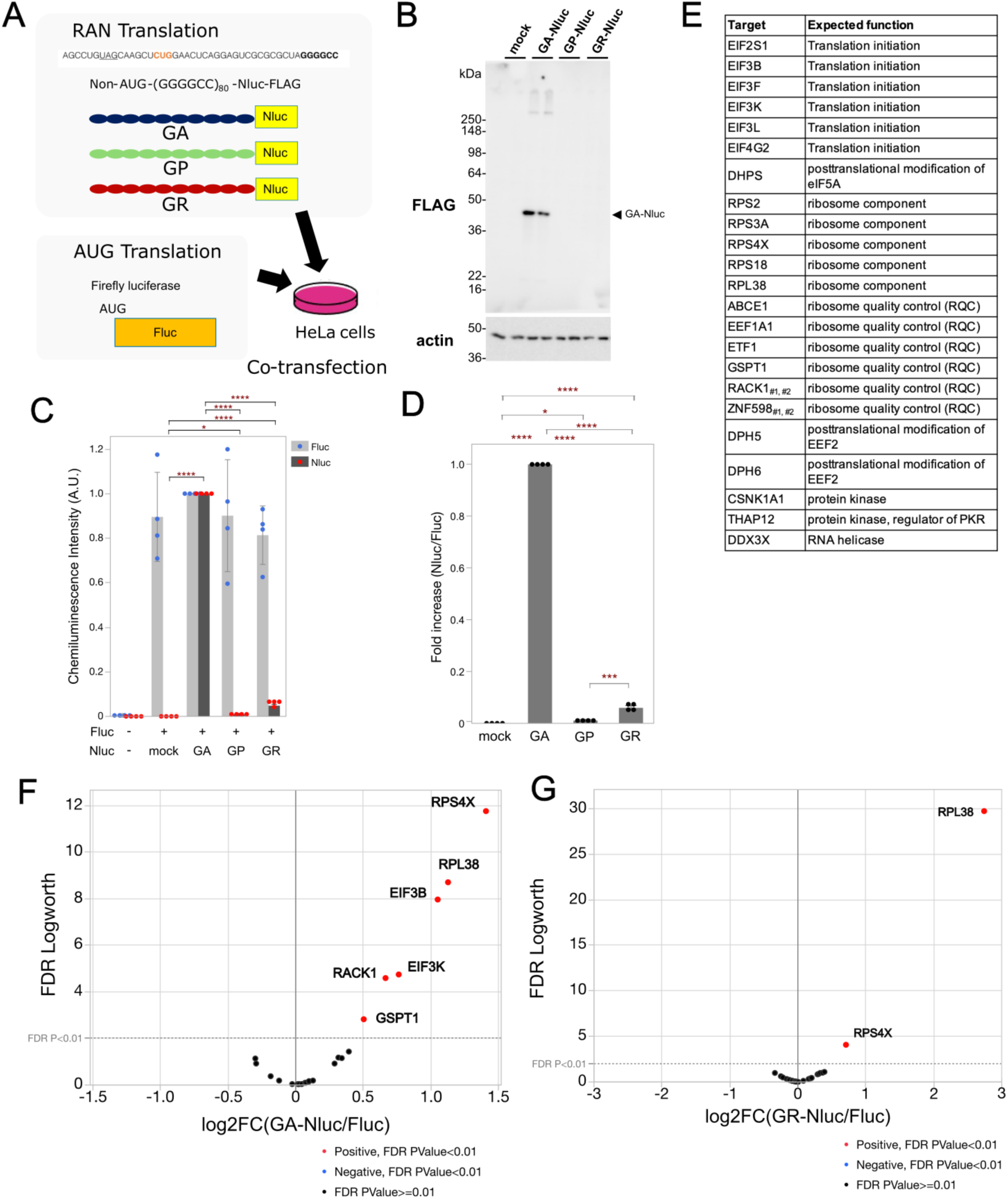
Dual-luciferase siRNA screen identifies candidate regulators of *C9ORF72* RAN translation. **(A)** Schematic of the dual-luciferase assay using Nanoluc-based reporters (Nluc) reflecting each reading frame of *C9ORF72* RAN translation and an AUG-driven Firefly luciferase reporter (Fluc) as a control for canonical translation. **(B)** Detection of RAN translation reporters by western blot. **(C)** Normalized chemiluminescence signals reflecting the relative expression levels of RAN-derived DPRs-Nluc and the AUG-Fluc reporter. n=4. **(D)** DPR luminescence normalized to AUG-Fluc, indicating the RAN/AUG translation ratio. **(C, D)** Graphs are shown in Mean ± SD. One-way ANOVA with Tukey-Kramer HSD post hoc test. *p<0.05, **p<0.01, ***p<0.001, ****p<0.0001. **(E)** List of target genes included in the siRNA screen and their expected functions. **(F)** siRNA screening results using the GA-Nluc reporter, showing log₂ fold changes (log2FC) of the GA-Nluc/AUG-Fluc ratio with corresponding FDR values (FDR Logworth). **(G)** Parallel siRNA screening results using the GR-Nluc reporter.

### Identification and characterization of RPL38 as a screening hit

Among the common top hits identified in both screens comparing AUG translation with poly-GA and poly-GR RAN translation, one notable candidate was RPL38. We therefore decided to focus our subsequent analyses on RPL38.

RPL38 is a component of the large (60S) ribosomal subunit and consists of 70 amino acid residues (Figure 2A). Among these residues, 14 are lysines (K) and 5 are arginine (R), making RPL38 a highly basic protein with an isoelectric point (pI) of 10.10. According to structural study (50) , RPL38 adopts a conformation composed of two α-helices and four β-sheet structures (Figure 2B) and directly binds to 28S ribosomal RNA of the large ribosome subunit. Within the ribosome, no direct interactions with other ribosomal proteins have been identified (Figure 2C). RPL38 is previously implicated in selective translation during vertebrate development, particularly in regulating the synthesis of homeobox transcription factors (51, 52) . In mouse, heterozygous knockout of *RPL38* is known as Tail short (Ts) phenotype showing pronounced homeotic transformation of the axial skeleton and homozygous knockout of *RPL38* is embryonic lethal (51, 53). According to the Protein Atlas database (https://www.proteinatlas.org/), the expression level of RPL38 in neurons of the normal human cerebral cortex increases with age (54) (Figure 2D).

**Figure 2.**
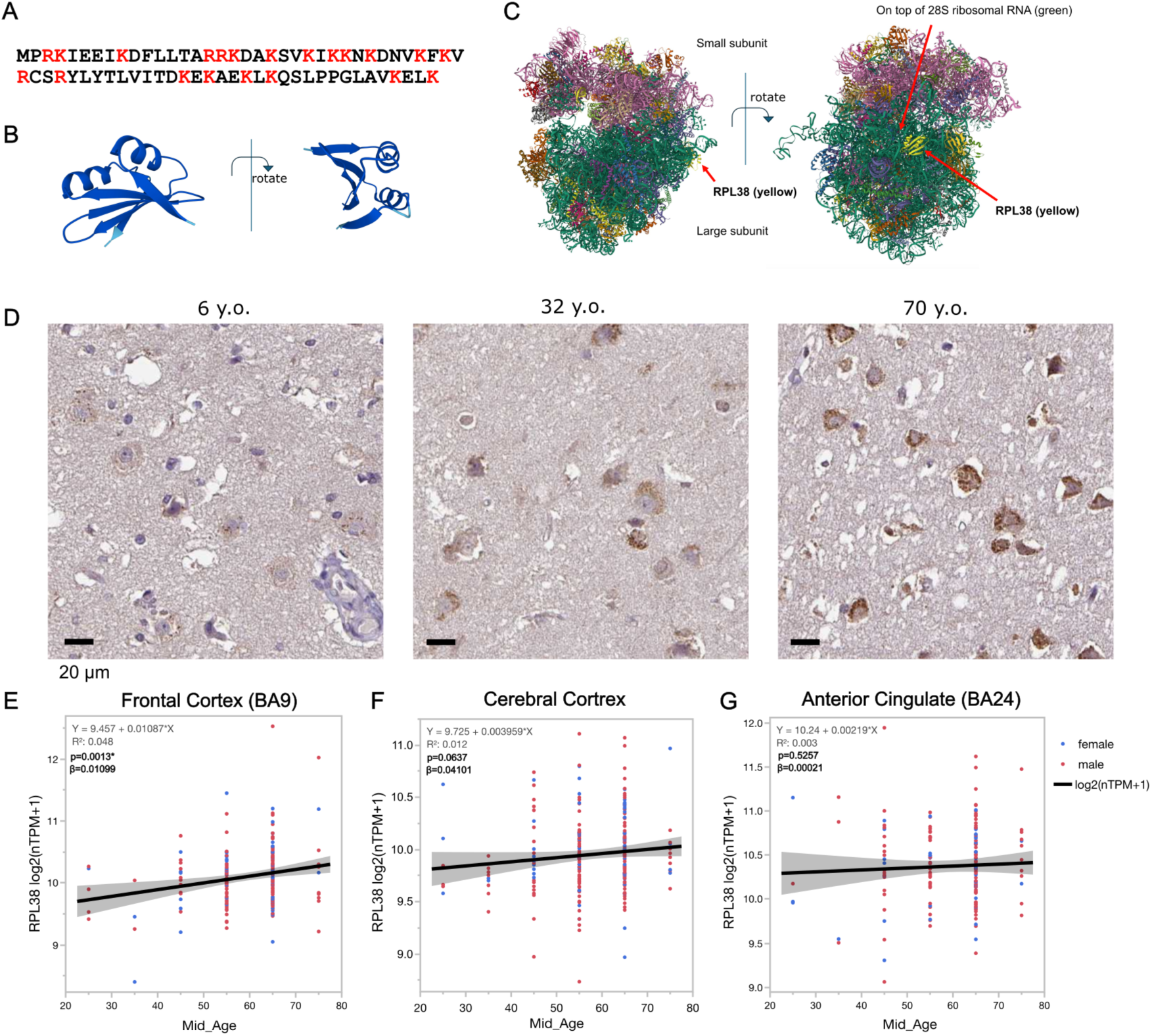
Structural and expression profiling of human RPL38. **(A)** Primary amino acid sequence (Basic amino acids are indicated in red) and **(B)** predicted tertiary structure of human RPL38. **(C)** Structural localization of RPL38 within the human ribosome (https://www.rcsb.org/3d-view/8QOI) (55, 56) . **(D)** RPL38 expression levels in neurons of the normal human cerebral cortex increase with age (the Protein Atlas database (https://www.proteinatlas.org)). **(E-G)** Relationship between the median age of samples (Mid_Age) and RPL38 expression levels in GTEx bulk RNA-seq data from the human frontal cortex **(E)**, cerebral cortex **(F)**, and anterior cingulate cortex **(G)**.

Reanalysis of bulk RNA-seq data from the GTEx dataset, accessed via the Human Protein Atlas, revealed a weak but statistically significant monotonic increase in RPL38 expression with age in the frontal cortex (BA9) (Spearman’s ρ = 0.1575, p = 0.0227). This age-associated trend remained consistent after adjustment for sex in a linear regression model using log2(nTPM + 1) values, although the effect size was modest (Figure 2E). In the cerebral cortex, a weak monotonic association with age was detected by Spearman correlation (ρ = 0.1342, p = 0.0322); however, this association did not remain statistically significant after adjustment for sex in linear regression analysis (Figure 2F). No significant age-related association was observed in the anterior cingulate cortex (BA24) (Spearman’s ρ = 0.0178, p = 0.8151) (Figure 2G).

Collectively, these findings highlight RPL38 as a structurally distinctive, basic component of the large ribosomal subunit, with modest and region-dependent age-associated expression changes in the human cerebral cortex.

### RPL38 knockdown preferentially suppresses canonical AUG translation

To validate the screening results and delineate the translational consequences of RPL38 depletion, we next analyzed its effects on AUG-dependent and RAN translation. Using independent siRNAs targeting different regions of RPL38, we confirmed that all three siRNAs, including the one used in the original screen, consistently increased RAN translation in the poly-GA and GP frames relative to AUG translation. Although one of the three siRNAs showed only a marginal effect in the poly-GR frame, the other two produced statistically significant increases, thus excluding potential off-target effect (Figure 3A). When we examined changes in the RAN and AUG translation reporters separately, we found that RPL38 knockdown had variable effects on RAN translation reporter expression depending on the reading frame or siRNA used, causing decreases, no change, or even modest increases (Figure 3B), whereas the AUG translation reporter showed a consistent and marked decrease in expression (Figure 3C). Together, these findings identify RPL38 as a ribosomal protein whose knockdown preferentially impairs canonical AUG translation, with comparatively minor effects on *C9ORF72* RAN translation.

**Figure 3.**
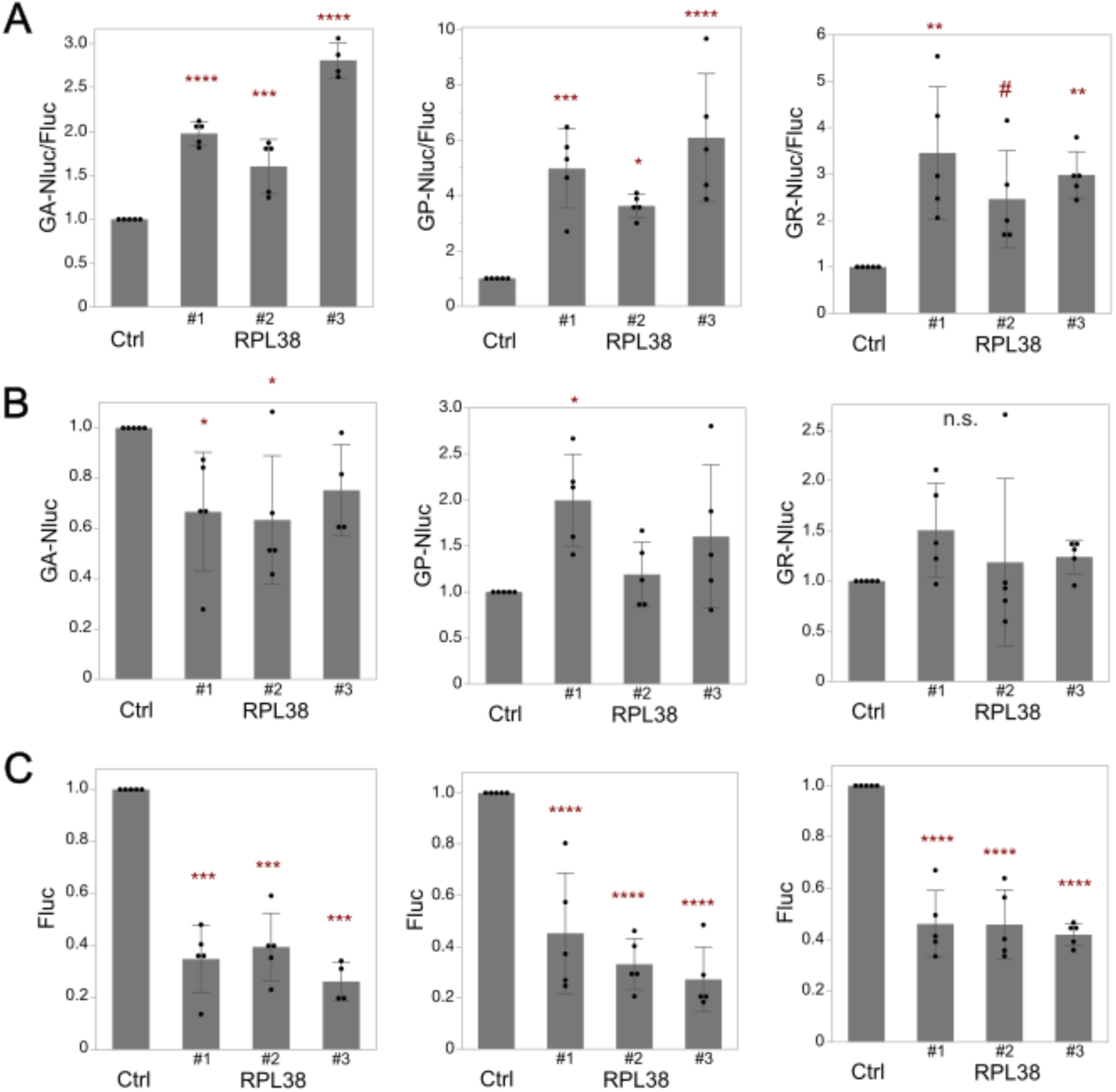
Validation of RPL38 knockdown effects on canonical AUG and *C9ORF72* RAN translation. **(A)** Normalized RAN (GA, GP, GR)/AUG reporter luminescence ratios following treatment with three independent RPL38-targeting siRNAs. **(B)** RAN (GA, GP, GR) Nluc luminescence ratios. **(C)** Corresponding luminescence signals of the AUG reporter under the same conditions. n=5, Each biological replicate is indicated by a dot. One-way ANOVA. All graphs are shown in Mean ± SD. One-way ANOVA with Dunnett’s post hoc test versus Ctrl knockdown. #p=0.0578, *p<0.05, **p<0.01, ***p<0.001, ****p<0.0001. n.s., not significant.

### RPL38 knockdown selectively impairs AUG-dependent translation and reduces global translational activity

To rule out the possibility that this result was an unintended artifact of the luciferase reporter system, we examined the expression of the poly-GA RAN translation product and AUG-derived EGFP by western blotting. This assay first confirmed efficient RPL38 knockdown by these siRNA (Figure 4A, D). Consistent with the dual-luciferase results, poly-GA levels did not exhibit a consistent and significant reduction (Figure 4A, B), whereas EGFP expression was consistently decreased (Figure 4D, E). In contrast, RT-qPCR analysis revealed no apparent changes in the expression levels of either G4C2 repeat RNA or EGFP mRNA following RPL38 knockdown further supporting RPL38 affect at translation step as predicted (Figure 4C, F). Initiation of translation at AUG codons constitutes the major portion of translational activity. If the expression level of RPL38 is reduced and thereby inhibits AUG-mediated translation, it is possible that the overall cellular translation level decreases. To evaluate the global translational activity following RPL38 knockdown, we performed a puromycin incorporation assay. Puromycin is an analogue of aminoacyl-tRNA that is incorporated into the C-terminus of nascent polypeptides during the elongation step of translation. After cell lysis, puromycin remains bound to nascent polypeptides and can be detected with an anti-puromycin antibody. Cells treated with cycloheximide (CHX), a translation inhibitor that blocks aminoacyl-tRNA translocation and thereby inhibits translation elongation, exhibited a significant decrease in puromycin incorporation in the cell lysates (Figure 4G, H). This result validated that the level of puromycin incorporation reflects the overall translational activity. Cells in which RPL38 was knocked down showed a significant reduction in puromycin incorporation compared with cells subjected to control knockdown using non-targeting siRNA, indicating that suppression of RPL38 expression leads to a global inhibition of translational activity (Figure 4G, H). These results suggest that RPL38 preferentially promotes AUG-initiated translation and plays an important role in maintaining global translational activity.

**Figure 4.**
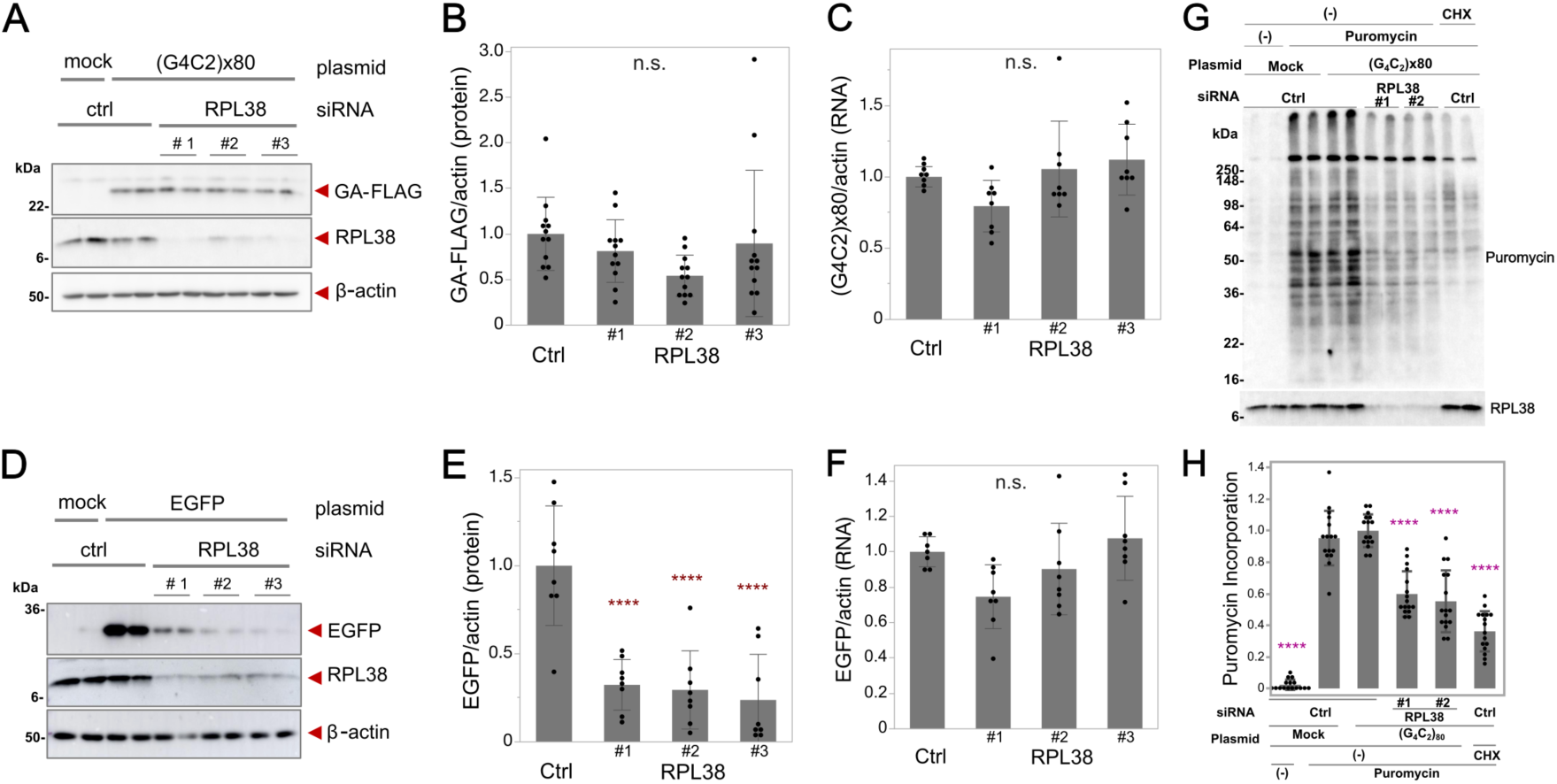
RPL38 knockdown impairs AUG-dependent and global translation. (A, **B)** RPL38 knockdown does not affect poly-GA expression derived from G4C2 repeat RAN translation. (n = 6 in biological duplicate). **(C)** G4C2 mRNA levels remain unchanged after RPL38 knockdown. (n = 4 in biological duplicate). **(D, E)** RPL38 knockdown significantly suppressed AUG-dependent EGFP expression in western blot. (n = 4, in biological duplicate). **(F)** EGFP mRNA levels were unchanged following RPL38 knockdown, as assessed by RT–qPCR. (n = 4, in biological duplicate). **(G)** Western blot analysis probed with anti-puromycin and anti-RPL38 antibodies. **(H)** Quantification of puromycin incorporation signal intensities (n=8, in biological duplicate). Graphs are presented as mean ± SD. Each biological replicate is indicated by a dot. Statistical analysis was performed using one-way ANOVA with Dunnett’s post hoc test versus control knockdown with G4C2 repeat or EGFP expression. *p<0.05, **p<0.01, ***p<0.001, ****p<0.0001, n.s., not significant.

### Dose-dependent effects of RPL38 knockdown on the RAN/AUG translation ratio

We next assessed how the extent of RPL38 knockdown quantitatively biases the balance between RAN and AUG translation. To address this, we transfected cells with siRNA at seven different concentrations ranging from 0.004 to 6.5 nM and examined the effects on RAN and AUG translation using a dual-luciferase assay. Regarding RAN translation, no changes were observed in the poly-GA or GP frames at any siRNA concentration, whereas the poly-GR frame showed a mild but statistically significant increase at higher siRNA concentrations (Figure 5A-C). In contrast, the AUG translation reporter exhibited a significant, dose-dependent decrease in reporter activity, beginning at relatively low siRNA concentrations (Figure 5D-F). Reflecting these results, RPL38 knockdown resulted in a dose-dependent increase in the RAN-to-AUG translation ratio (Figure 5G-I). A linear, dose-dependent increase in the RAN-to-AUG translation ratio upon RPL38 knockdown indicates that the effect is quantitatively dependent on RPL38 expression level. Collectively, these findings show that the balance between RAN and AUG translation is quantitatively tuned by RPL38 levels, with progressive RPL38 depletion selectively impairing AUG-dependent translation and thereby increasing the RAN-to-AUG ratio.

**Figure 5.**
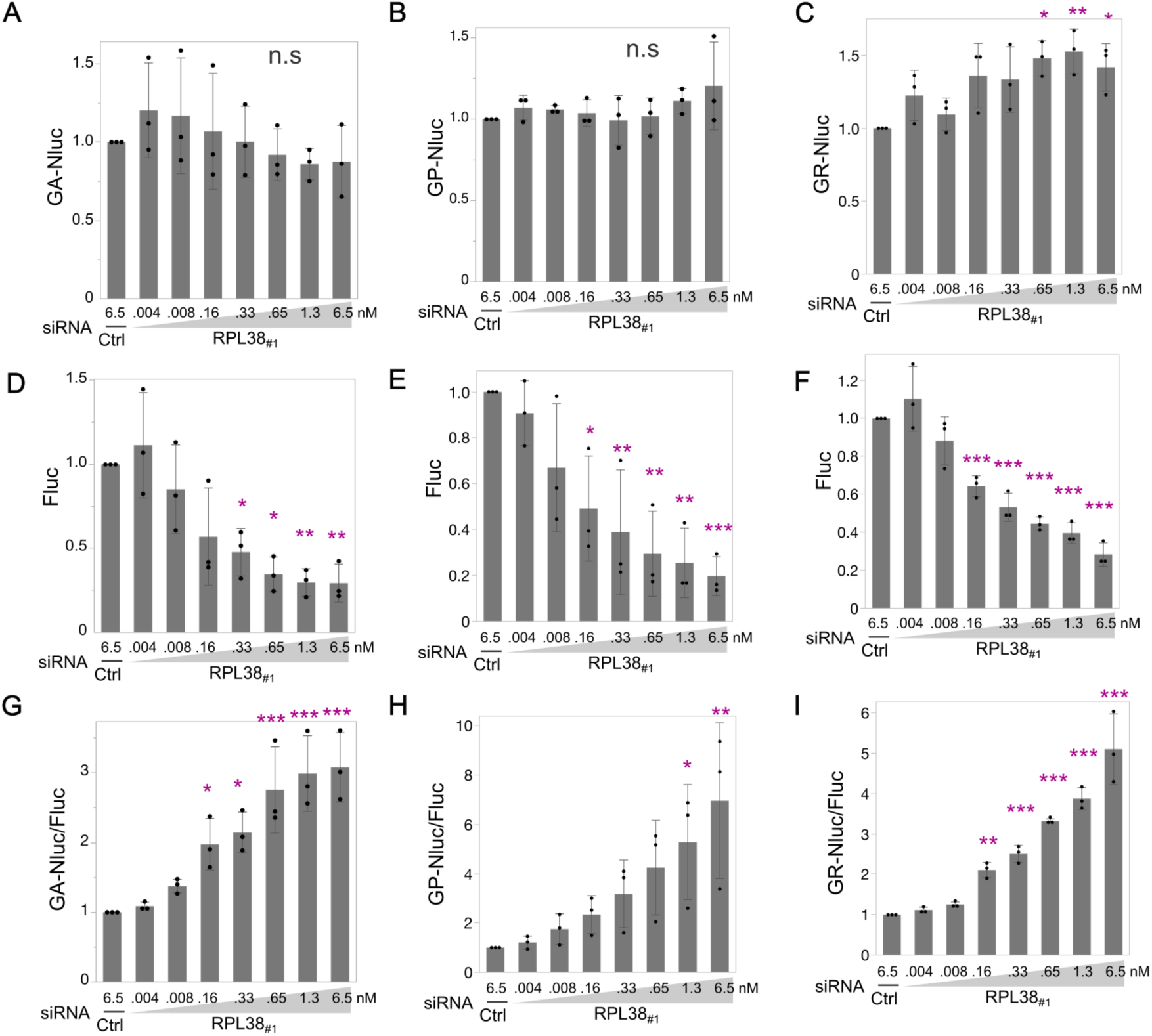
Graded RPL38 knockdown differentially affects RAN and AUG-dependent translation. Dose-dependent effects of graded RPL38 knockdown on RAN translation and canonical AUG-dependent translation using dual luciferase reporter system. (**A-C)** RAN (GA, GP, GR) Nluc luminescence ratios. **(D-F)** Corresponding Fluc luminescence signals of the AUG reporter, **(G-I)** Normalized RAN (GA, GP, GR)/AUG reporter luminescence ratios. Mean ± SD (n = 3). One-way ANOVA with Dunnett’s post hoc test versus Ctrl KD. *p<0.05, **p<0.01, ***p<0.001, ****p<0.0001. n.s., not significant.

### Conversion of CUG to AUG renders poly-GA translation sensitive to RPL38

The above results revealed that reduced expression of RPL38 had a strong impact on AUG-initiated translation. The RAN translation of poly-GA initiates from a near-cognate codon, CUG, located 24 nucleotides upstream of the GGGGCC repeat. If this CUG is replaced with the canonical start codon AUG, would the change in poly-GA translation levels be influenced by the expression level of RPL38? To investigate this point, we performed the dual luciferase experiment using a plasmid in which the CUG codon was replaced with an AUG codon (Figure 6A) (29) . In contrast to the CUG-initiated poly-GA RAN translation reporter used in Figure 5A, the AUG-initiated poly-GA translation reporter exhibited a dose-dependent decrease in reporter activity in response to siRNA targeting RPL38 (Figure 6B). The AUG-initiated Fluc reporter showed a similar result (Figure 6C). Consistent with these findings, the ratio of AUG-GA-Nluc to Fluc did not change with the concentration of siRNA against RPL38 (Figure 6D). Furthermore, in western blot analysis using three different siRNAs targeting RPL38, the AUG-initiated poly-GA showed a significant decrease upon RPL38 knockdown (Figure 6E, F), in contrast to the CUG-initiated poly-GA RAN translation product where no change in poly-GA expression level was observed following RPL38 knockdown (Figure 4A, B). Together, these results demonstrate that replacing the near-cognate CUG start codon with a canonical AUG converts poly-GA translation into an RPL38-dependent, AUG-initiated process, thereby rendering DPR production sensitive to RPL38 expression levels.

**Figure 6.**
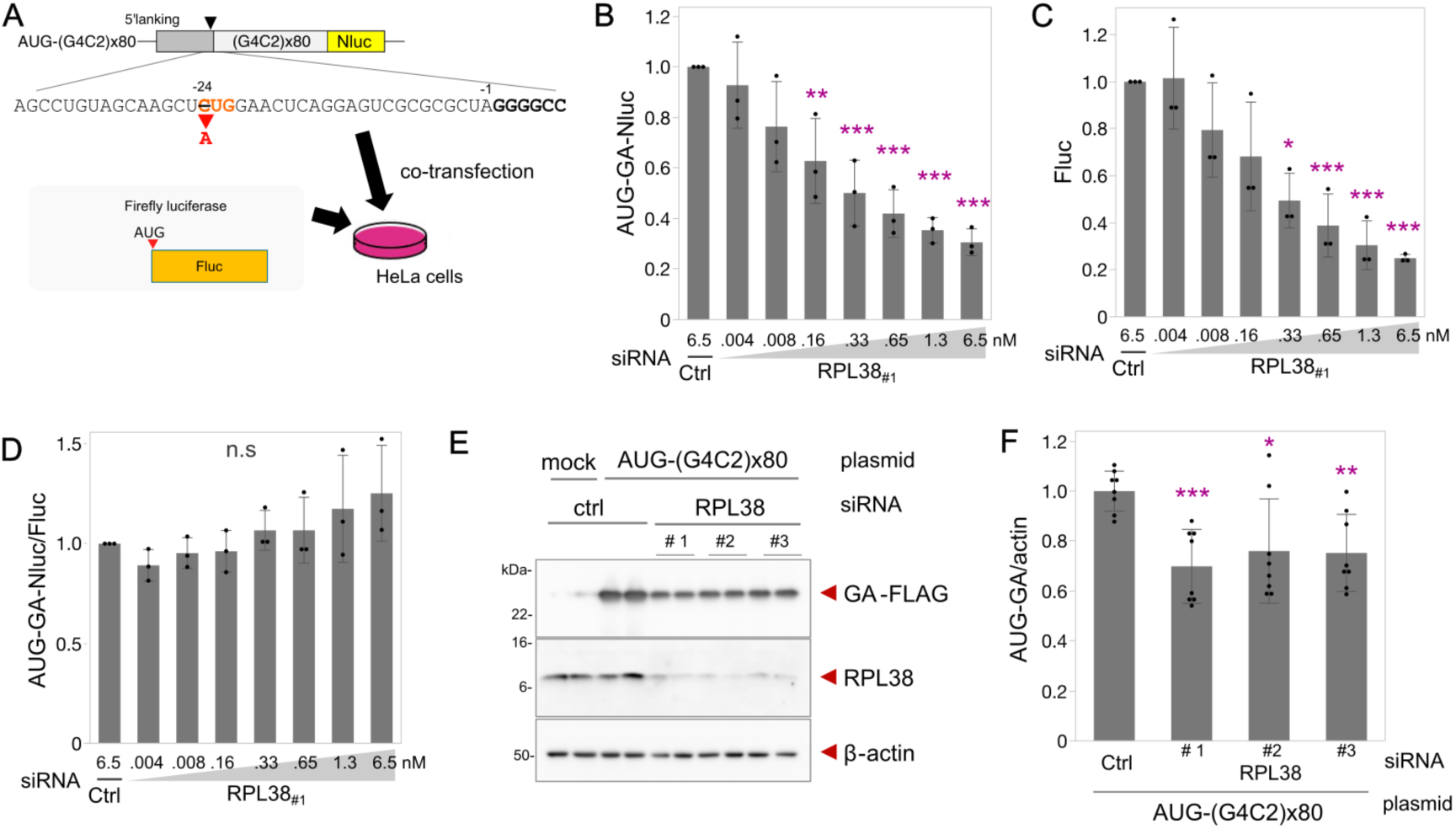
AUG initiation renders poly-GA translation sensitive to RPL38 depletion. **(A)** Schematic of the dual-luciferase reporter assay using a (G4C2)x80 CUG-to-AUG mutant construct with Nluc in the poly-GA reading frame. **(B-D)** Dose-dependent effects of graded RPL38 knockdown on AUG-dependent poly-GA translation (B) and canonical AUG-dependent Fluc translation **(C)**. **(D)** AUG-GA-Nluc/AUG-Fluc. (n = 3). **(E, F)** RPL38 knockdown significantly suppressed AUG-dependent poly-GA expression in western blot (n = 4, in biological duplicate). All graphs are shown in mean ± SD. One-way ANOVA with Dunnett’s post hoc test versus Ctrl KD. *p<0.05, **p<0.01, ***p<0.001, ****p<0.0001. n.s., not significant.

### RPL38 depletion selectively reduces large ribosomal subunit abundance

To investigate how reduced RPL38 expression influences ribosomal composition within cells expressing G4C2 repeat, we conducted polysome profiling. Cytoplasmic fractions, obtained from HeLa cells after removal of the nuclear fraction, were subjected to sucrose density gradient centrifugation, and absorbance at 260 nm was monitored across the fractions. In cells expressing the G4C2 repeat, RPL38 knockdown led to a marked reduction in the 60S/40S ratio, reflecting the balance between large and small ribosomal subunits (Figure 7A, B) compared with non-targeting control knockdown cells. A260 profile suggest the level of 40S is relatively constant but the level of 60S is reduced (Figure 7A). Similarly, RPL38 knockdown led to a marked reduction in the 80S/40S ratio, representing the proportion of monosomes relative to small subunits (Figure 7A, C). These findings indicate that depletion of RPL38 results in a relative deficiency of large ribosomal subunits. In contrast, the 80S/60S ratio, which reflects the efficiency of large subunit association with small subunits to form monosomes, remained unchanged upon RPL38 knockdown, suggesting that diminished RPL38 expression does not impair the docking efficiency between ribosomal subunits (Figure 7A, D). Together, these results demonstrate that RPL38 is required for maintaining the proper balance between ribosomal subunits, and that its depletion selectively reduces large subunit abundance without affecting subunit joining efficiency.

**Figure 7.**
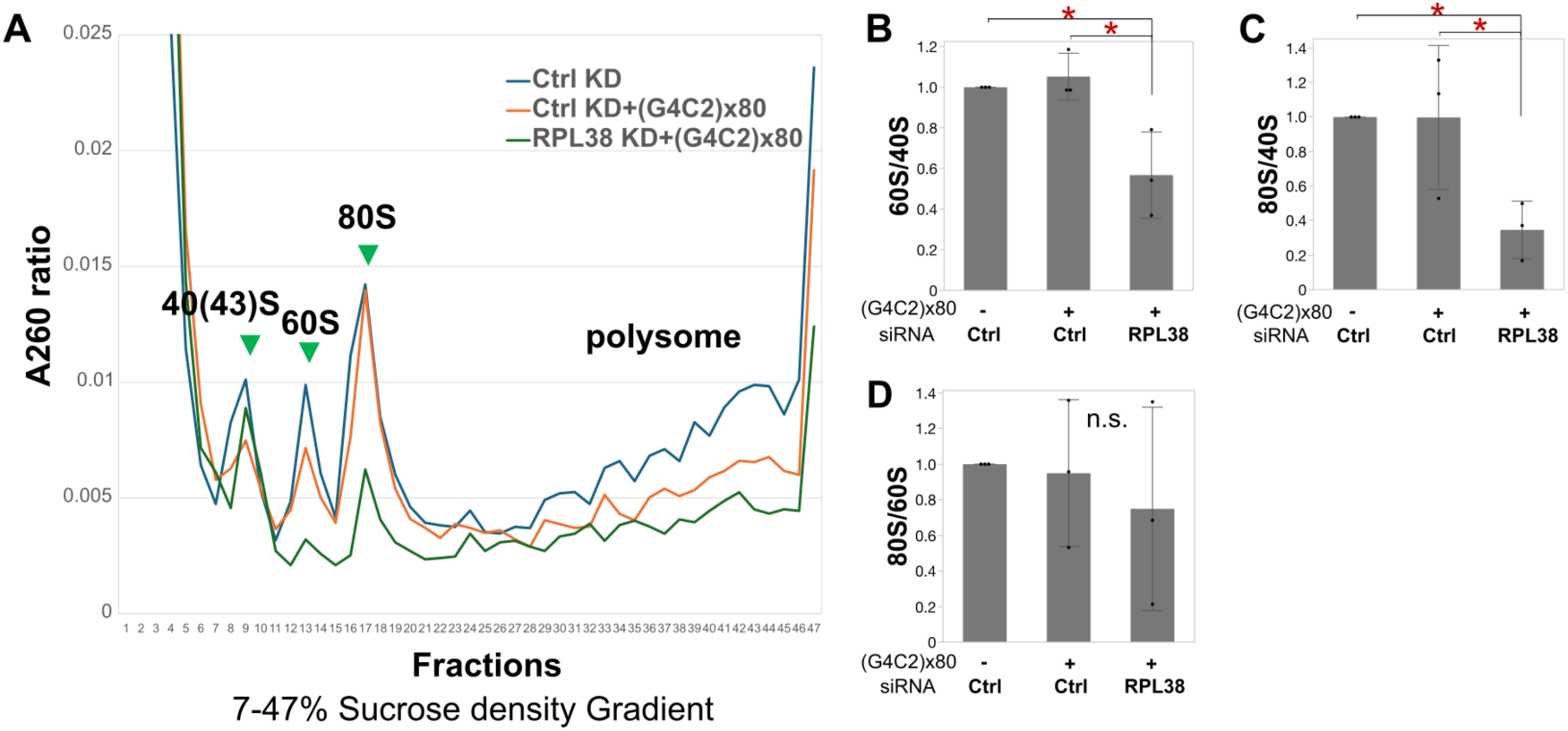
Polysome profiling reveals a reduction in the large ribosomal subunit upon RPL38 depletion. **(A)** Polysome profiles of HeLa cells expressing (G4C2)80 repeats with or without RPL38 knockdown. **(B)** A260 ratio of the 60S large subunit to the 40S small subunit. **(C)** A260 ratio of the 80S monosome to the 40S small subunit. **(D)** A260 ratio of the 80S monosome to the 60S large subunit. Graphs are presented as mean ± SD (n = 3). Statistical analysis was performed using one-way ANOVA with Tukey’s post hoc test **(A-D)**. *p<0.05. n.s., not significant.

### RPL38 depletion induces frame-specific remodeling of *C9ORF72* RAN translation

We next asked whether an imbalance between ribosomal subunits caused by reduction of RPL38 could differentially affect individual reading frames of *C9ORF72* RAN translation. To address this, we reanalyzed the results of our siRNA screen in which the relative contribution of RAN and AUG-initiated translation was quantified using poly-GA and poly-GR RAN translation reporters (Fig. 1). Specifically, for each knockdown condition, we calculated the ratio of the GR-Nluc/AUG-Fluc value to the GA-Nluc/AUG-Fluc value. Strikingly, knockdown of RPL38 or CSNK1A1 resulted in a significant increase in this ratio, whereas knockdown of EIF3B, EIF4X, or EIF3K led to a significant decrease (Figure 8A). To further validate that reduction of RPL38 exerts differential effects on individual RAN translation reading frames, we newly constructed a GA-Fluc reporter and co-transfected it together with the GR-Nluc reporter into cells in which RPL38 was knocked down to varying extents. This experimental design enabled a direct, side-by-side comparison of frame-specific differences in *C9ORF72* RAN translation under graded levels of RPL38 depletion. A consistent, dose-dependent increase in the poly-GR/poly-GA ratio was detected using multiple independent siRNAs against RPL38 (Figure 8 B-D). These results demonstrate that RPL38 exerts frame-specific effects on *C9ORF72* RAN translation, indicating that perturbation of ribosome composition differentially modulates individual RAN reading frames.

**Figure 8.**
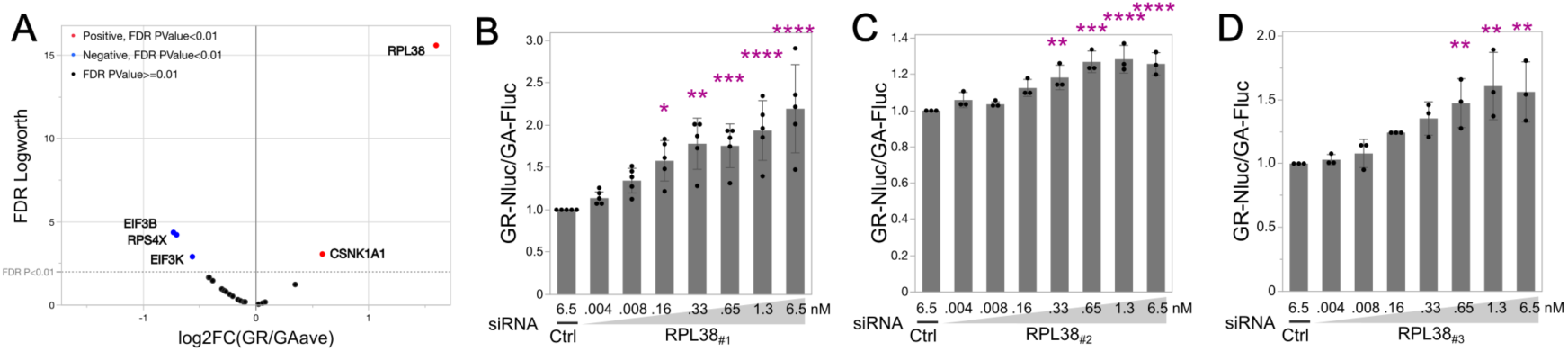
RPL38 depletion alters frame selection of *C9ORF72* RAN translation. **(A)** Reanalysis of the siRNA screen showing the ratio of the GR-Nluc/AUG-Fluc value to the GA-Nluc/AUG-Fluc value, displayed as a volcano plot. **(B-D)** Dose-dependent effects of graded RPL38 knockdown on poly-GR RAN translation (Nluc) and poly-GA RAN translation (Fluc), respectively, assessed using a dual-luciferase reporter system. **(B)** siRNA for RPL38#1 (n=5), **(C)** siRNA for RPL38#2 (n=3), and **(D)** siRNA for RPL38#3 (n = 3). Mean ± SD. One-way ANOVA with Dunnett’s post hoc test versus Ctrl KD. *p<0.05, **p<0.01, ***p<0.001, ****p<0.0001.

Lastly, to evaluate *in vivo* effect of RPL38 knockdown on *C9ORF72* RAN translation, we examined the previously established *Drosophila* disease model which express 44 G4C2 repeats under the eye-specific GMR-Gal4 driver (57) . This model contains endogenous 5′flanking leader sequence adjacent to *C9orf72* G4C2 repeats including the near-cognate CUG start codon. It also contains a 3′ GFP tag in the GR reading frame, which enables fluorescence imaging and quantitation of poly-GR expression levels. Knockdown of endogenous RPL38 significantly enhanced poly-GR expressions in the fly eyes compared to control knockdown (Fig. 9, A and B). On the other hand, its knockdown reduces AUG-dependent EGFP expression in the control EGFP fly (Fig. 9, C and D). Thus, consistent with the results of the cellular models, RPL38 also differently regulates RAN translation and AUG-dependent translation in a *Drosophila* model of *C9ORF72*-FTD/ALS.

**Figure 9.**
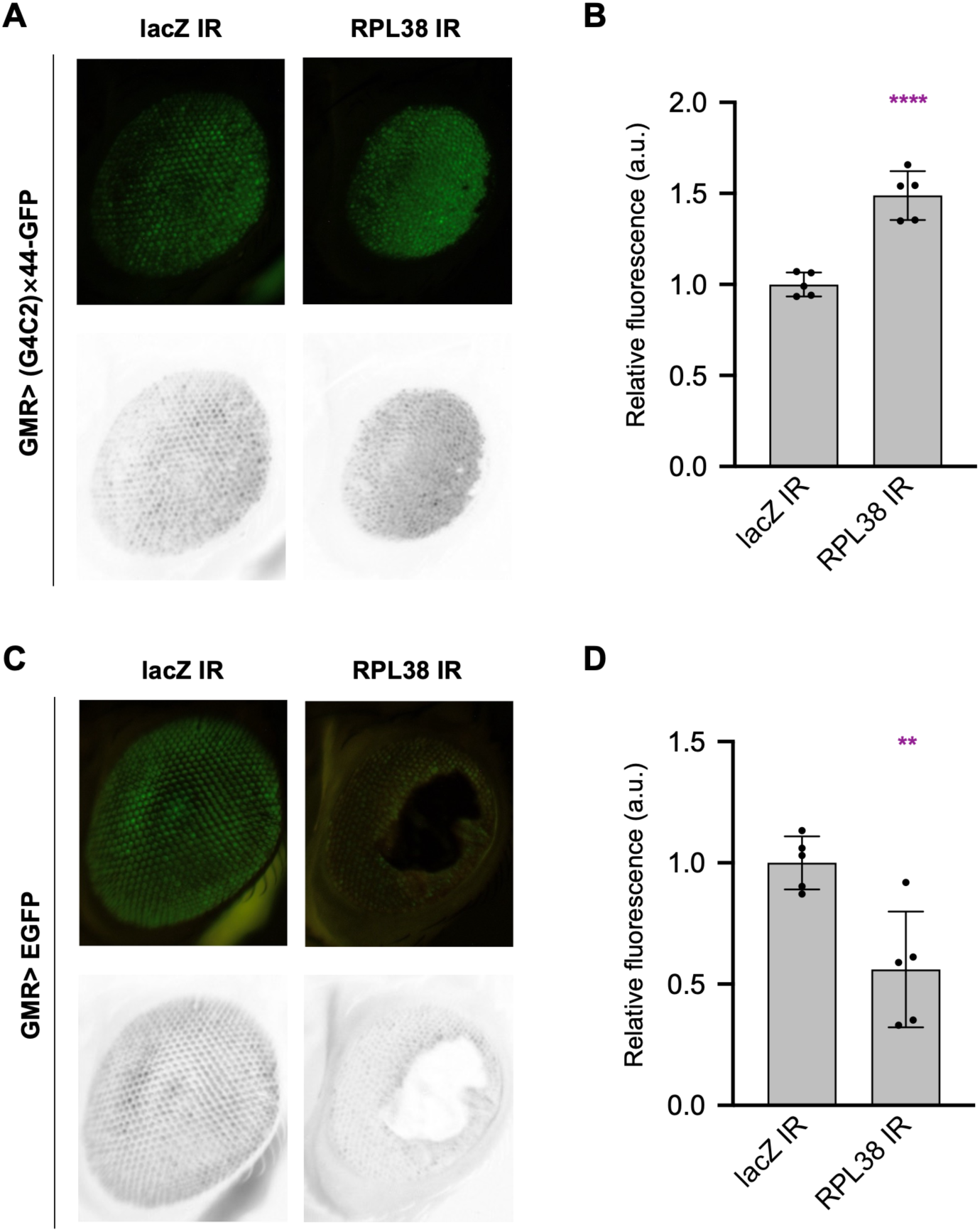
RPL38 knockdown enhances GR-frame RAN translation but suppresses AUG-dependent EGFP expression in the *Drosophila* eye. **(A)** Representative images of GFP fluorescence in *Drosophila* eyes expressing a 44-repeat GR-frame reporter with a C-terminal GFP tag and the 5′ leader sequence. Upper panels show raw GFP fluorescence images and lower panels show grayscale images of the green channel. RPL38 knockdown increased GFP fluorescence intensity. **(B)** Quantification of GFP fluorescence intensity per unit eye area in the genotypes shown in (A). n = 5 per genotype. Statistical significance was assessed using a bilateral unpaired t-test. **(C)** Representative images of GFP fluorescence in control flies expressing EGFP in the eye by canonical AUG-initiated translation. Upper panels show raw GFP fluorescence images and lower panels show grayscale images of the green channel. RPL38 knockdown reduced GFP fluorescence intensity. Fluorescence-negative regions in the center of RPL38-knockdown eyes correspond to necrotic tissue. Because RPL38 knockdown caused marked toxicity, fluorescence analysis was performed in regions excluding necrotic areas. **(D)** Quantification of GFP fluorescence intensity per unit eye area in the genotypes shown in (C). n = 5 per genotype. Statistical significance was assessed using a bilateral unpaired t-test. Graphs are shown as mean ± SD. **p<0.01, ****p<0.0001.

In summary, dual-luciferase knockdown screens identified RPL38 as a key factor that preferentially supports canonical AUG-initiated translation while exerting multilayered effects on *C9ORF72* RAN translation. RPL38 depletion reduced large ribosomal subunit abundance and globally suppressed translation, yet selectively increased the RAN-to-AUG translation ratio. Importantly, graded knockdown of RPL38 revealed a dose-dependent, frame-specific increase in the poly-GR-to-poly-GA RAN translation ratio, demonstrating that ribosomal subunit imbalance differentially modulates individual RAN reading frames. Consistent with these observations, RPL38 knockdown in the *Drosophila* eye increased GR-frame RAN translation while reducing AUG-dependent EGFP expression, extending this translational bias to an *in vivo* context.

## Discussion

RPL38 was originally identified as a determinant of ribosome-mediated translational specificity, selectively promoting the translation of a subset of Hox mRNAs without broadly suppressing global protein synthesis (51). However, loss of individual ribosomal proteins is generally associated with reduced ribosome abundance and global translational repression (58–60). In this context, our findings extend the role of RPL38 beyond transcript-specific regulation and identify it as a regulator of start-codon selection and translational output.

We show that depletion of RPL38 preferentially suppresses canonical AUG-initiated translation while relatively preserving initiation at near-cognate codons, thereby shifting the balance between canonical and non-canonical translation. This effect is quantitatively dependent on RPL38 levels and is accompanied by a selective reduction in 60S ribosomal subunit abundance without impairment of subunit joining efficiency, consistent with a defect in large subunit biogenesis. Previous studies have highlighted the importance of ribosomal composition in translational control (45, 46), and our results further suggest that 60S subunit availability acts as a limiting factor that enforces stringent AUG selection during translation initiation.

Our findings also reveal that ribosomal subunit imbalance differentially affects translation across reading frames. Prior work has shown that insufficiency of 40S ribosomal proteins, including RPS25 and RPS26, suppresses non-canonical translation in repeat expansion disorders (49, 61), indicating that small subunit integrity is required for efficient RAN translation. In contrast, reduction of the 60S subunit protein RPL38 does not uniformly inhibit non-canonical initiation but instead alters frame usage, favoring the GR reading frame over GA. These observations suggest that the 40S and 60S subunits play distinct and potentially opposing roles in regulating initiation stringency and frame selection. Consistent with the cell-based findings, RPL38 knockdown in the *Drosophila* eye enhanced GR-frame RAN translation while suppressing canonical AUG-dependent translation, supporting the physiological relevance of this translational bias *in vivo*.

Mechanistically, a relative deficiency of the large ribosomal subunit is expected to limit productive 80S assembly, thereby preferentially suppressing efficient AUG-dependent initiation. Under these conditions, the availability of preinitiation complexes and small ribosomal subunits may exceed that of the large subunit, reducing the stringency of start-codon selection. As a result, suboptimal start sites, including near-cognate codons, are more likely to be recognized as initiation sites. Thus, reduced initiation stringency in RPL38-depleted cells may allow non-canonical translation to be maintained despite global translational repression (Figure 10).

**Figure 10.**
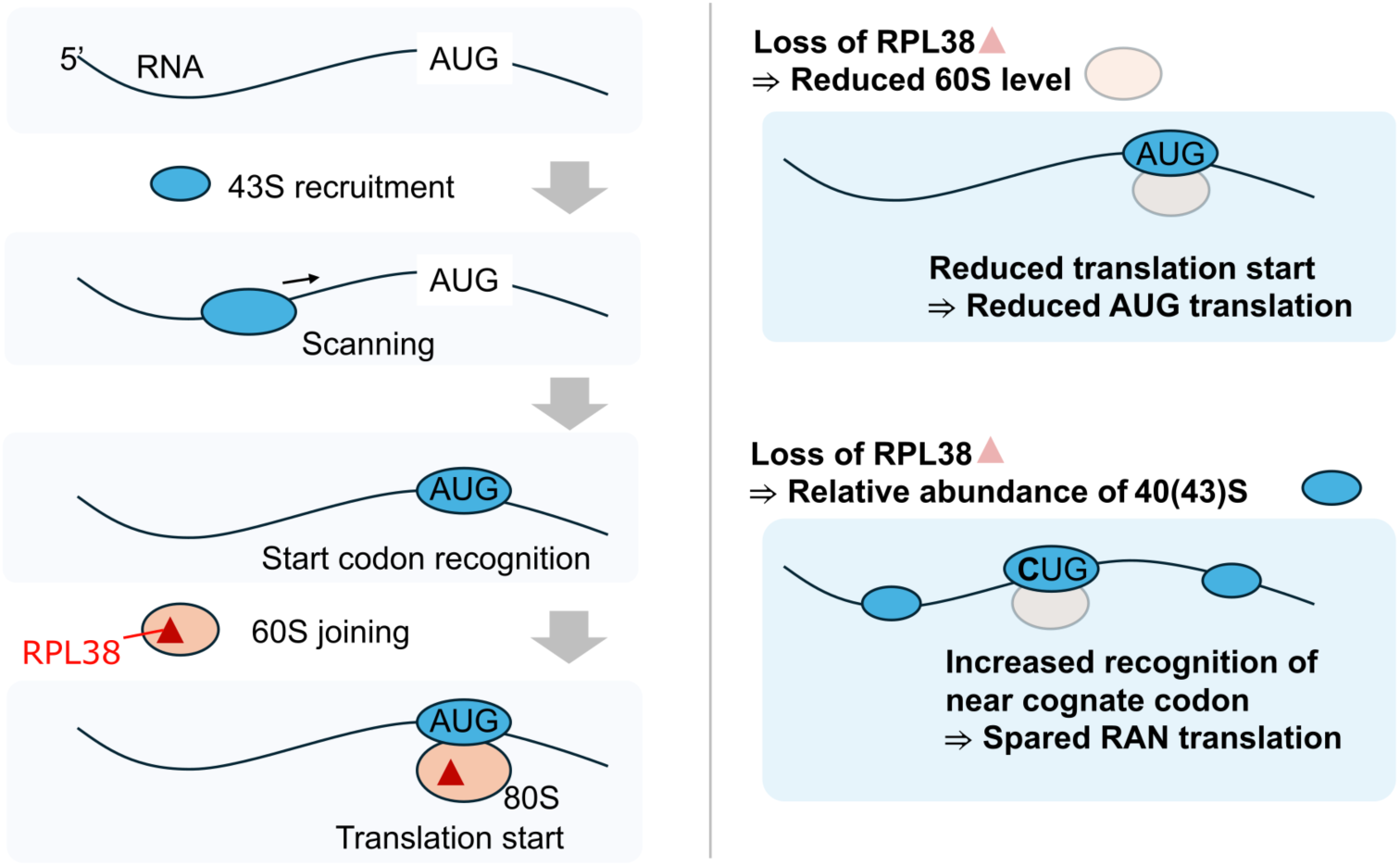
Model for how reduced 60S ribosomal subunit availability modulates start-codon selection and frame usage during *C9ORF72* RAN translation. (Left) Under normal conditions, the 43S preinitiation complex scans mRNAs from the 5′ end and efficiently recognizes optimal AUG start codons, followed by recruitment of the 60S large subunit to form the 80S ribosome and initiate translation. (Upper right) Upon RPL38 depletion, reduced availability of the 60S subunit limits productive 80S assembly, thereby preferentially suppressing efficient AUG-dependent initiation. (Lower right) Under these conditions, an increased relative abundance of scanning preinitiation complexes and small ribosomal subunits may reduce the stringency of start-codon selection, allowing initiation at suboptimal near-cognate codons. This shift may preserve non-canonical RAN translation despite global translational repression and alter frame usage within repeat RNA. This model is based on the observed reduction in 60S subunit abundance, differential sensitivity of AUG- and CUG-initiated translation to RPL38 depletion, and dose-dependent changes in RAN-to-AUG translation and frame-specific output.

Within C9ORF72 repeat RNA, initiation efficiency differs between reading frames. The poly-GA frame utilizes a CUG start codon and exhibits relatively higher initiation efficiency, whereas poly-GR initiation is less efficient (28). Under conditions of reduced initiation stringency, this difference may shift frame usage toward less efficient initiation events, leading to an increased poly-GR/poly-GA ratio. Thus, ribosomal subunit imbalance can act not only on start-codon selection but also on reading frame usage.

Importantly, our findings likely reflect the consequences of impaired 60S subunit biogenesis rather than the function of mature heterogeneous ribosomes selectively lacking RPL38. RPL38 depletion induces a global reduction in large subunit abundance, indicating that the observed effects arise from altered ribosome homeostasis at the level of subunit availability. This interpretation is consistent with studies showing that perturbation of ribosome biogenesis broadly affects translational capacity (58–60).

Taken together, our study shows that selective perturbation of 60S ribosomal subunit homeostasis by RPL38 depletion reshapes the balance between canonical and non-canonical translation in a frame-dependent manner. These findings establish ribosomal subunit availability as a previously underappreciated determinant of start-codon stringency, frame selection, and C9ORF72 RAN translation dynamics, and provide a mechanistic basis for how ribosome homeostasis shapes pathological non-canonical translation in a disease-relevant context.

## Materials and Methods

### Cell culture

HeLa cells were cultured in DMEM containing 10% FCS and penicillin 100 U/ml/ and streptomycin 100 μg/ml.

### Plasmids

The plasmids were driven by either the CMV promoter and expressed the endogenous 113-nucleotide 5′ leader sequence upstream of the G4C2 repeat, followed by 80 G4C2 repeats and corresponding peptide tags in each reading frame (22) . The 5′ leader sequence contains a near-cognate CTG codon located 24–22 nucleotides upstream of the first G of the G4C2 repeat, which has been proposed to serve as the initiation codon for poly-GA translation. Substitutions of CTG to CCG within the 5′ leader sequence were introduced by oligonucleotide ligation (29).

Nanoluc luciferase (Nluc)- or Firefly luciferase (Fluc)-based RAN translation reporters for poly-GA, poly-GP, and poly-GR were constructed by fusing each luciferase with FLAG-tag to the C-terminus of the corresponding dipeptide repeat in each reading frame. The DNA fragment encoding EGFP was amplified by PCR and subcloned into the BamHI/NotI sites of the pcDNA3.1 hygro (+) vector.

### Antibodies & reagent

The following antibodies were used at the indicated dilutions: anti-β-actin ((SIGMA #A5316) 1/1000 or (Cell Signaling #4970) 1/1000), anti-DYKDDDDK (FLAG) Tag (Cell Signaling #2368S) 1/1000, anti-RPL38 (Bethyl A305-412A) 1/2000, anti-GFP (B2, Santa Cruz, sc-9996) 1/500, anti-puromycin, 12D10 (MERCK #MABE343) 1/25000, anti-poly-GA (proteintech #24492-1-AP) 1/1000 or (Millipore #MABN889) 1/1000. The following reagents were used: puromycin (INVIVOGEN), cycloheximide (CHX) (Nakarai tesque 06741-91, CAS 66-81-9), Halt Protease and Phosphatase Single-Use Inhibitor Cocktail(100X) (Thermo 78442), and Anti-Rabbit IgG (H+L), HRP Conjugate (Promega, W401B), Anti-mouse IgG (H+L), HRP Conjugate (Promega, W402B).

### siRNA-mediated knockdown and plasmid transfection

In knockdown experiments, HeLa cells cultured in 24-well plates and 5 pmol of siRNA/well of 24-well plates (10 nM) unless otherwise indicated were reverse-transfected with RNAiMAX reagent (Invitrogen) and incubated overnight. In the overexpression experiments, repeat plasmids or mock plasmids were transfected using the lipofectamine LTX reagent (Invitrogen). Six hours after transfection, the medium was replaced with fresh medium. The following day, cells were washed with PBS and used for subsequent analyses. The siRNA sequences used for knockdown experiments were listed in supplementary table.

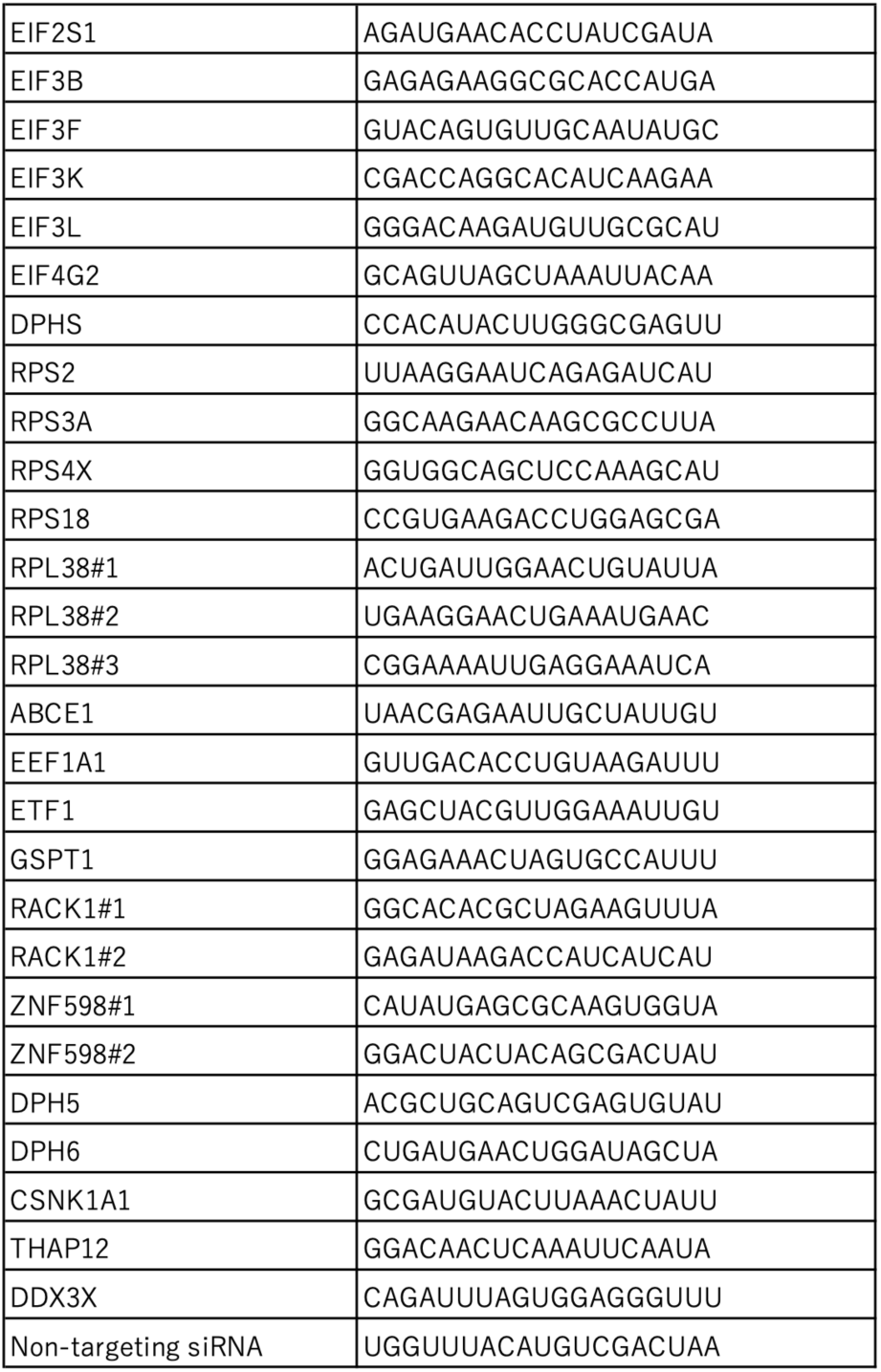
Supplementary Table -List of siRNA-

### Dual-Luciferase Reporter Assay

Dual-luciferase assay was performed by using Nano-Glo Dual-Luciferase Reporter Assay System (N1630, Promega) according to manufactures protocol. pGL4.53 [luc2/PGK] (E5011, Promega) was used as AUG-Fluc reporter. To identify knockdown conditions with significant effects compared to the control (Ctrl), we applied the Response Screening platform, which fits one-way linear models for each condition in JMP Student Edition 18.2.2 (SAS Institute Inc., Cary, NC, USA). For each comparison, raw p-values were calculated and adjusted for multiple testing using the Benjamini–Hochberg false discovery rate (FDR) method. Effect sizes were expressed as log2 fold changes (log2FC) relative to the control group. The control mean value was used as the baseline for fold change calculations, and log2 transformation was applied prior to analysis. Significance thresholds were set at FDR-adjusted p < 0.01 and an absolute log2FC > 0.5. For visualization, volcano plots were generated using the Graph Builder in JMP, plotting log2FC (x-axis) against LogWorth (y-axis, defined as −log10(p)). Reference lines were drawn at log2FC = ±0.5 and LogWorth = 2 (corresponding to p = 0.01).

### Correlation analysis of RNAseq data with age

Gene expression levels were obtained from publicly available RNA-seq datasets and were quantified as normalized transcripts per million (nTPM) as provided by the data source (https://www.proteinatlas.org/ENSG00000172809-RPL38/tissue/cerebral+cortex#imid_15090338, accessed at Dec, 23, 2025). No additional normalization was applied. To reduce right-skewness inherent to RNA-seq expression data and to facilitate interpretation, nTPM values were log2-transformed after adding a pseudocount of 1 log2(nTPM+1) prior to statistical analysis. Age information was available in 10-year categories (e.g., 20–29, 30–39 years). Each age category was converted to its midpoint value and treated as a continuous variable in subsequent analyses. To assess monotonic associations between age and gene expression without assuming linearity, Spearman’s rank correlation coefficient (ρ) was calculated between age and log2(nTPM+1) values. Two-sided p-values were reported.

To estimate the magnitude and direction of age-related changes in gene expression while adjusting for sex, we fitted linear regression models using the standard least squares method: log2(nTPM+1)=𝛽0+𝛽1⋅Age+𝛽2⋅Sex+𝜀 where Age represents the midpoint of each age category and Sex was included as a categorical covariate. Regression coefficients (β), 95% confidence intervals, p-values, and coefficients of determination (R²) were obtained.

Sex-adjusted regression lines and 95% confidence intervals were visualized using predicted values derived from the fitted regression models. All statistical tests were two-sided, and p-values < 0.05 were considered statistically significant.

### Western blotting

Cells were dissolved in a RIPA buffer supplemented with a Protease Inhibitor Cocktail (SIGMA) with EDTA and sonicated using a Bioruptor II. The samples were boiled for 10 min in the presence of 1×SDS loading buffer. The samples were loaded onto a 10% to 15% Tris-Glycine gel. After SDS-PAGE, samples were transferred onto PVDF or Nitrocellulose (pore size 0.2 μm for RPL38) membranes (Merck). After blocking with I-block (Thermo) for 1 hour, the membrane was incubated overnight with a diluted primary antibody. The following day, the membrane was washed 3 times with TBST for 5 minutes and incubated with an HRP-labelled secondary antibody for 60 minutes. The membrane was then washed three times with TBST for 10 min. Chemiluminescent signals were detected using an Amersham Imager 680 (GE Healthcare). Signals were quantified using Image J.

### RT-qPCR

Total RNA was prepared using the RNeasy and Qiashredder kit (Qiagen). RNA preparations were treated with Turbo DNA-free kit (Thermo Fisher Scientific) to minimize residual DNA contamination. Two micrograms of RNA were used for reverse transcription with M-MLV Reverse Transcriptase (Promega) using oligo-(dT) 12-18 primer (Invitrogen). RT-qPCR was performed using the QuantStudio 6 Real-Time PCR System (Applied Biosystems) with TaqMan technology. Primers and probes were designed (IDT) for 3′ TAG region of repeat constructs (repeat TAG primer) and EGFP. Repeat TAG, Primer 1: TCT CAA ACT GGG ATG CGT AC, Primer 2: GTA GTC AAG CGT AGT CTG GG, Probe/56-FAM/TG CAG ATA T/Zen/C CAG CAC AGT GGC G/3IABkFQ/(38). EGFP, Primer 1: GCA CAA GCT GGA GTA CAA CTA, Primer 2: TGT TGT GGC GGA TCT TGA A, Probe /56-FAM/AG CAG AAG A/Zen/A CGG CATCAA GGT GA/3IABkFQ/(38). Primer/ probe sets for Human ACTB (β-actin), Hs.PT.39a.22214847 (IDT) were used as endogenous control. Each sample was paired with no reverse transcription controls showing < 1/2^10 (ΔCT > 10) signal when compared to reverse transcribed samples, thus excluding contamination of plasmid DNA-derived signal. Each biological sample was analyzed in technical duplicate or triplicate. Signals of repeat RNA were normalized to β-actin according to the ΔΔCT method (44).

### Puromycin incorporation assay

Puromycin incorporation assay (alternatively called SUnSET assay) was performed according to Schmidt et al (62) . HeLa cells were knocked down with siRNAs for RPL38 or non-targeting control. Next day, those cells were transfected with repeat or mock plasmids. A subset of cells were additionally treated with 20 μM of CHX for 20 min as a control for translational inhibition. Additional treatment with 10 μg/ml puromycin (INVIVOGEN) for 10 min for pulse labelling of ongoing translation. Cells were washed once with PBS then served for western blotting with anti-puromycin antibody (44) .

### Polysome profiling assay

Polysome profiling analysis according to Pringle et al. (44, 63) was performed with modification. HeLa cells reverse transfected overnight in the presence or absence of RPL38 or control siRNA (30 pmol/10cm dish) were further transfected with or without 80 repeats of G_4_C_2_ repeat transfection and then treated with 100 μg/ml CHX in PBS for 3 min at 37 °C to halt elongation. Cells were immediately washed, collected, lysed in low-salt lysis buffer [20 mM Tris-HCl pH7.4, 50 mM KCl, 10 mM MgCl_2_, 1% Triton X-100, 0.5%(w/v) sodium deoxycholate, 1 mM 1, 4-dithiothreitol (DTT), 1x Halt Protease and Phosphatase Inhibitor, EDTA free (Thermo, 78442), 100 μg/ ml CHX] in the presence of RiboLock RNase inhibitor (Thermo, EO0381) on ice for 10 min. Nuclei were removed by centrifugation at 2,000 x g for 5min. 500 μl of cytoplasmic fraction was then loaded on the top of the 7%-47% linear sucrose gradient in low-salt buffer [20 mM Tris-HCl pH7.4, 50 mM KCl, 10 mM MgCl_2_, 100 μg/ ml CHX] formed by gradient master (Biocomp) in a open top centrifuge tube (7030 SETON Scientific). Then ultracentrifugation was performed at 260,808 x g (=39,000 rpm on the SW41 rotor, XPN-80 ultracentrifuge (Beckman Coulter)) with slow acceleration up to top speed, 90 min centrifugation at top speed, maximum deceleration until the rotor reaches 2,000 rpm and then no brakes until the rotor automatically stops. After the ultracentrifugation, 250 μl of fractions were sequentially isolated from the top of the gradient by piston gradient fractionator (Biocomp). Since continuous UV monitoring system was not available in our facility, each fraction was measured OD_260_ (A260) to monitor polysome profiles.

### Fly stocks

All fly stocks were maintained and crossed at 25 °C on standard cornmeal-yeast-glucose medium. Transgenic flies carrying the GMR-Gal4, UAS-lacZ IR, and UAS-LDS-(G4C2)×44-GFP transgenes have been described previously (57, 64, 65) . Transgenic flies carrying the UAS-EGFP and UAS-RPL38 IR were obtained from the Bloomington *Drosophila* Stock Center (#6874) and the Vienna *Drosophila* Resource Center (#28565) (66) , respectively.

### External fly eye fluorescent imaging

Female flies expressing the UAS-LDS-(G4C2) × 44-GFP or UAS-EGFP under the control of the eye-specific GMR-Gal4 driver were crossed with male flies carrying UAS-RPL38 IR or UAS-lacZ IR. Heads from 1-2-day-old progeny were dissected, and fluorescence images of compound eyes were acquired using a stereomicroscope SZX10 equipped with the digital camera DP23 (Evident Scientific) with a GFP filter. All images were acquired under the same exposure conditions. Raw GFP images were converted to greyscale, and the mean grey values of compound eyes were measured using ImageJ. Areas of eye necrosis were excluded from the analysis.

## Statistical analysis

Statistical analyses were performed using JMP Student Edition 18.2.2 (SAS Institute Inc., Cary, NC, USA).

## Data availability

No large-scale datasets are associated with this study.

## Acknowledgements

We thank Dr. Yuya Kawabe (Minoh Neuropsychiatric Sanatorium), and Ms. Miyabi Arai (The University of Osaka) for helpful discussions; Ms. Ayako Goto (Center for Medical Research and Education, Graduate School of Medicine, The University of Osaka) for assistance with polysome profiling assays. KMo is supported by JST FOREST Program under grant number JPMJFR200Z, the JSPS KAKENHI grant numbers JP20H05927 (KMo, YN), JP22K19492; JP24K02382, JP25K22600, JP25K02592 (MI, KMo), AMED grant number JP23dk0207066; SENSHIN Medical Research Foundation (KMo, TM), The Mochida Memorial Foundation for Medical and Pharmaceutical Research and Takeda Science Foundation. SG is supported by JSPS KAKENHI JP24K23408 and JP25K19078, ST by 22K07559, TM by JP25K23931. RU by Grant-in-Aid for JSPS Research Fellows JP25KJ1747. RU, KMi and YA by Next Generation Researcher Development Project by The University of Osaka. MI by AMED grant number JP23bm0804034. Stocks obtained from the Bloomington Drosophila Stock Center (NIH P40OD018537) and the Vienna Drosophila Resource Center were used in this study.

## Author contributions

KMo conceived and supervised the project, analyzed the data, performed statistical analyses, prepared the figures and tables, and wrote the manuscript with input from all the authors. SG generated key repeat constructs and performed polysome profiling. SK performed most experiments under the supervision of KMo. YF and YN conducted and analyzed the *Drosophila* experiments and prepared the corresponding figure. RU established the dual-luciferase assay. SG, YF, RU, KMi, YA, ST, SA, TM, and YN provided conceptual input and contributed to data interpretation. KMo and MI coordinated the study. All authors approved the final manuscript.

## Abbreviations

The abbreviations used are:

ALS: amyotrophic lateral sclerosis
A260/OD260: absorbance/optical density at 260 nm
CHX: cycloheximide
DMEM: Dulbecco’s modified Eagle’s medium
DPR(s): dipeptide repeat protein(s)
DTT: dithiothreitol
EGFP: enhanced green fluorescent protein
eIF: eukaryotic initiation factor
FCS: fetal calf serum
Fluc: Firefly luciferase
FTD: frontotemporal dementia
G4C2: GGGGCC repeat
GTEx: Genotype-Tissue Expression
HRP: horseradish peroxidase
IRES: internal ribosome entry site
ISR: integrated stress response
KD: knockdown
log2FC: log2 fold change
Nluc: NanoLuc luciferase
nTPM: normalized transcripts per million
pI: isoelectric point
PBS: phosphate-buffered saline
PKR: double-stranded RNA-activated protein kinase (EIF2AK2)
RAN: repeat-associated non-AUG
rRNA: ribosomal RNA
RNA-seq: RNA sequencing
RT–qPCR: reverse transcription quantitative PCR
SD: standard deviation
SDS-PAGE: sodium dodecyl sulfate–polyacrylamide gel electrophoresis
siRNA: small interfering RNA
SUnSET: surface sensing of translation
TBST: Tris-buffered saline with Tween 20
uORF: upstream open reading frame.

